# Dectin-1/CARD9-induction of the TFEB and TFE3 gene network is dispensable for phagocyte anti-*Aspergillus* activity in the lung

**DOI:** 10.1101/2023.06.13.544785

**Authors:** Mariano A. Aufiero, Neta Shlezinger, Mergim Gjonbalaj, Kathleen A.M. Mills, Andrea Ballabio, Tobias M. Hohl

## Abstract

Myeloid phagocytes of the respiratory immune system, such as neutrophils, monocytes, and alveolar macrophages, are essential for immunity to *Aspergillus fumigatus*, the most common etiologic agent of mold pneumonia worldwide. Following engulfment of *A. fumigatus* conidia, fusion of the phagosome with the lysosome, is a critical process for killing conidia. TFEB and TFE3 are transcription factors that regulate lysosomal biogenesis under stress and are activated by inflammatory stimuli in macrophages, but it is unknown whether TFEB and TFE3 contribute to anti-*Aspergillus* immunity during infection. We found that lung neutrophils express TFEB and TFE3, and their target genes were upregulated during *A. fumigatus* lung infection. Additionally, *A. fumigatus* infection induced nuclear accumulation of TFEB and TFE3 in macrophages in a process regulated by Dectin-1 and CARD9 signaling. Genetic deletion of *Tfeb* and *Tfe3* impaired macrophage killing of *A. fumigatus* conidia. However, in a murine immune competent *Aspergillus* infection model with genetic deficiency of *Tfeb* and *Tfe3* in hematopoietic cells, we surprisingly found that lung myeloid phagocytes had no defects in conidial phagocytosis or killing. Loss of TFEB and TFE3 did not impact murine survival or clearance of *A. fumigatus* from the lungs. Our findings indicate that myeloid phagocytes activate TFEB and TFE3 in response to *A. fumigatus*, and while this pathway promotes macrophage fungicidal activity *in vitro*, genetic loss can be functionally compensated at the portal of infection in the lung, resulting in no measurable defect in fungal control and host survival.

## INTRODUCTION

*Aspergillus fumigatus* is a ubiquitous opportunistic mold that is the primary cause of invasive aspergillosis. *A. fumigatus* was recently classified by the World Health Organization as a pathogen of critical priority (1). Individuals with hematologic malignancies or those receiving high doses of corticosteroids often have quantitative and/or qualitative defects in innate immune function that render them susceptible to invasive disease upon inhalation of *A. fumigatus* conidia, the airborne infectious propagules (2). In the lungs of patients with defective innate immune function, conidia can germinate and form tissue-invasive hyphae that cause tissue injury and can result in mortality (3, 4). Due to challenges in early diagnosis, limited numbers of antifungal drug classes, and emerging resistance to front-line agents (5), overall mortality rates for patients with invasive aspergillosis remain high, with fungus-attributable rates of 20-30% in high-risk groups (6). Thus, a better understanding of how the respiratory immune system responds to *Aspergillus* infection may lead to novel therapeutic strategies to improve patient outcomes.

Clearance of *Aspergillus* from the lungs is primarily mediated by innate immune phagocytes, including resident alveolar macrophages, recruited neutrophils, inflammatory monocytes, and monocyte-derived dendritic cells (Mo-DCs), all of which phagocytose and kill *Aspergillus* conidia before they can germinate in the lung (7, 8). Following phagocytosis, the phagosome undergoes a maturation process via sequential fusion steps with components of the endocytic compartment, the final step of which is fusion with the lysosome, the primary degradative compartment of the cell (9). This maturation process and fusion with the lysosome allows for the acquisition of various antimicrobial factors and properties such as an acidified pH (via the vacuolar ATPase), the assembly of NADPH oxidase which produces antimicrobial reactive-oxygen species (ROS), and the activation of lysosomal hydrolases which degrade phagosomal contents (10). Blocking these effector functions, such as by blocking phagosomal acidification with bafilomycin or abrogating ROS production by mutations in NADPH oxidase components, attenuates the ability of innate immune cells to kill *Aspergillus in vitro* and *in vivo* (11–13).

The requirement for lysosomes in the clearance of phagocytic targets has long been appreciated, but it has only recently been understood how the lysosome can be dynamically regulated in cells. In the past decade, the transcription factors TFEB and TFE3 have been functionally characterized as master regulators of lysosomal biogenesis and autophagy in response to cellular stress (14–17). Various stimuli can lead to TFEB and TFE3 activation, including starvation, lysosomal damage, unfolded proteins, and infection (14). Activation of these transcription factors leads to lysosomal gene transcription and lysosomal biogenesis to enable the cell to cope with stress conditions. Bacterial pathogens or the fungal pathogen, *Candida albicans,* can activate these transcription factors in macrophages and promote downstream effector functions (18–20). In macrophages, inflammatory stimuli such as LPS or bacteria promote lysosomal gene expression and the degradative capacity of lysosomes in a TFEB- and TFE3-dependent manner (18, 19). In turn, TFEB- and TFE3-activated lysosomes promote macrophage effector functions that enhance bacterial clearance (18). TFEB-regulated expression of *Irg1* and itaconate production contribute to macrophage antibacterial functions (21). TFEB and TFE3 can also directly regulate the expression of pro-inflammatory cytokines in cultured macrophages (19, 22). Thus, TFEB and TFE3 support macrophage effector functions in response to bacterial stimuli, but the *in vivo* role of TFEB and TFE3 during bacterial and fungal infection in mammalian hosts remains largely undefined, with no studies on microbial burden in infected organs or on host survival. In this study, we observed that *A. fumigatus* drives TFEB target gene transcription in infected neutrophils *in vivo*, and nuclear localization of TFEB in macrophages *in vitro*. TFEB and TFE3 were required for optimal macrophage killing of *A. fumigatus in vitro* but were ultimately dispensable for lung leukocyte killing of *A. fumigatus* and survival during *A. fumigatus* respiratory infection in otherwise immune competent mice.

## RESULTS

### Lysosomal genes are upregulated in lung neutrophils during *A. fumigatus* infection

We hypothesized that fungal engagement by myeloid phagocytes leads to the upregulation of genetic pathways that are important for their fungicidal activity. To identify which pathways are upregulated upon fungal engagement in the lung, we challenged mice with Alexa Fluor 633 (AF633)-labeled *A. fumigatus* conidia via the intratracheal route, isolated bystander (AF633^-^) and fungus-engaged (AF633^+^) neutrophils from the lung by FACS 24 hours post-infection (hpi), and performed RNA-seq on sorted cells. Using KEGG pathway analysis, we compared genes that were upregulated in fungus-engaged neutrophils compared to bystander neutrophils. We found that genes associated with lysosomes were significantly overrepresented among all upregulated genes in fungus-engaged neutrophils (Fig. 1A).

**Figure 1:**
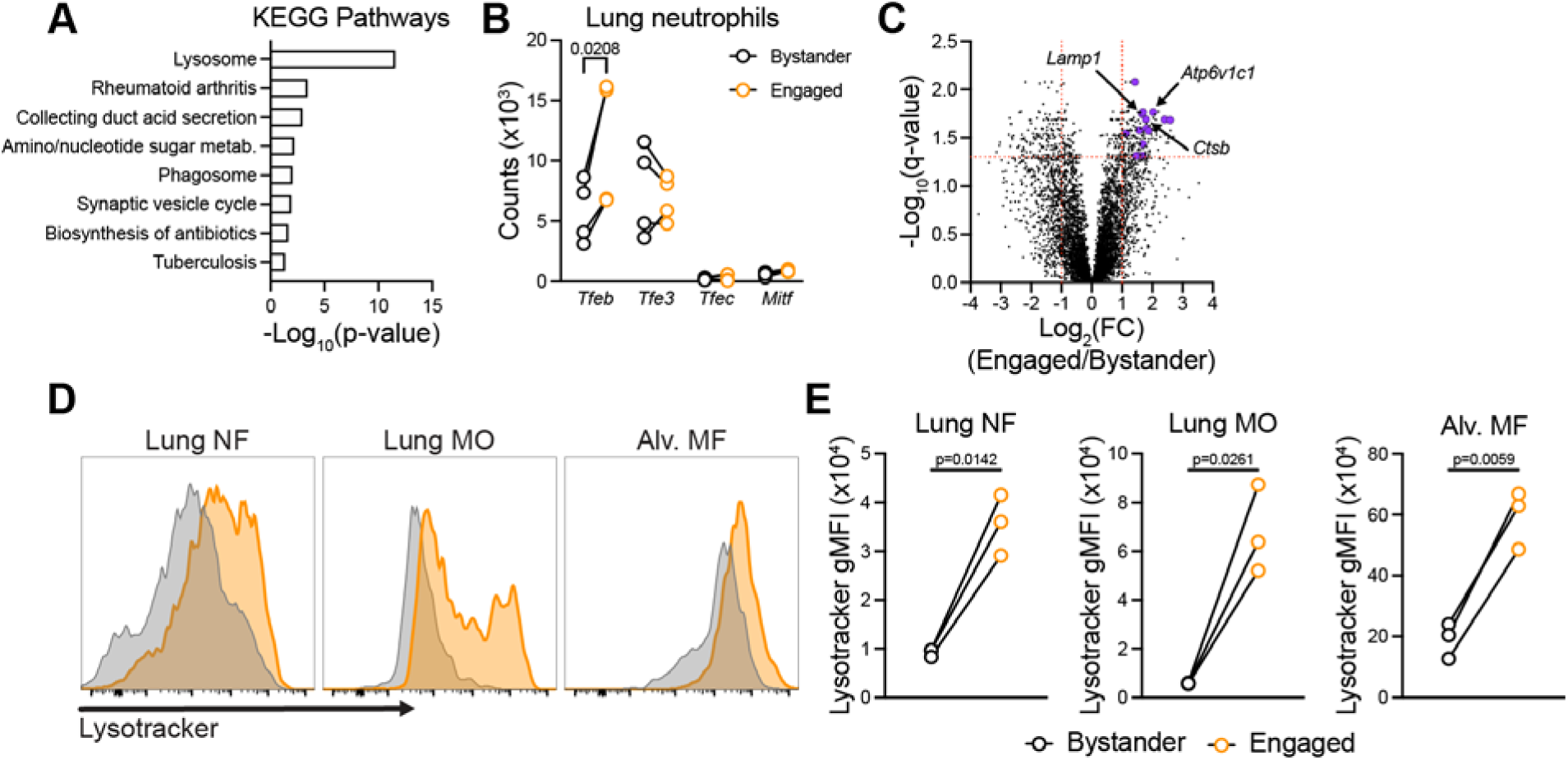
Fungus-engaged neutrophils upregulate lysosome genes, and lung phagocytes expand their lysosomal compartment in response to fungal engagement. **(A)** Bar chart displaying the statistically significant KEGG pathways of upregulated genes in engaged (AF633^+^) neutrophils relative to bystander (AF633^-^) neutrophils sorted from the lungs of *A. fumigatus* infected mice 24 hours post infection. **(B)** Volcano plot showing differential gene expression in engaged neutrophils relative to bystander neutrophils. Significantly upregulated TFEB target genes are colored purple. Vertical dashed lines indicate log_2_ fold change of −1 or 1. Horizontal line indicates q-value = 0.05. **(C)** Scatter dot plots indicating the gene counts for MiTF/TFE transcription factor family members. Dots represent independent experiments and lines link data from the same experimental replicate. **(D)** Representative histograms that display fluorescence intensity of LysoTracker green in the indicated lung leukocytes populations 24 hpi with with AF633-labeled *Aspergillus*. Gray histogram indicates bystander phagocytes and the orange histogram indicate engaged phagocytes. (A, B, and C) Data are pooled from 4 independent experiments. (D) Data are representative of 2 independent experiments. Statistics: (C) Two-way RM ANOVA with Sidak’s multiple comparisons test. (D) Paired t-test.

The transcription factors TFEB and TFE3 control lysosomal gene expression in a variety of physiological contexts (14, 16, 23). We found that neutrophils express *Tfeb* and *Tfe3*, but not the other MiTF/TFE family members, *Mitf* and *Tfec* (Fig. 1B). In addition, *Tfeb* is upregulated upon fungal engagement (Fig. 1C), consistent with a known feed-forward loop in TFEB activity (24). When compared to a published list of confirmed TFEB target genes (25), fifteen confirmed TFEB target genes were significantly upregulated (see supplemental table 1 for list of genes) in fungus-engaged neutrophils, and no TFEB target genes were significantly downregulated (Fig. 1C).

In addition to lung neutrophils, alveolar macrophages, monocytes, and monocyte-derived dendritic cells (Mo-DCs) phagocytose and kill conidia during *A. fumigatus* infection (7, 11). To assess whether fungal engagement led to an expansion of lysosomes in lung phagocytes, we challenged C57BL/6J (WT) mice with AF633-labeled conidia, and stained lung cells *ex vivo* with LysoTracker, a fluorescent dye that labels lysosomes, at 24 hpi. Fungus-engaged neutrophils had increased LysoTracker signal compared to bystander neutrophils, consistent with an expansion of their lysosomal compartment (Fig. 1D). Additionally, we observed a similar increase in the LysoTracker signal in engaged lung monocytes and engaged alveolar macrophages compared to bystander cells. These data indicate that myeloid phagocytes expand their lysosomal compartment when cells engulf *Aspergillus* conidia in the lung.

### *A. fumigatus* conidia activate TFEB and TFE3 in macrophages in part through C-type lectin receptors

Bacteria and *C. albicans* can activate TFEB and TFE3 in macrophages (19, 20, 22), though the mechanism by which fungal recognition is coupled to TFEB and TFE3 activation has not been defined. To test whether *A. fumigatus* could activate TFEB and TFE3, we challenged immortalized bone marrow derived macrophages (iBMDMs) with *A. fumigatus* heat-killed swollen conidia and assessed nuclear translocation of endogenous TFEB and TFE3 by confocal microscopy, as previously described (19). We observed rapid TFEB and TFE3 nuclear localization following *A. fumigatus* infection, which peaked as early as 1 hpi for TFE3 and at 4 hpi for TFEB. The nuclear to cytoplasmic ratio of TFEB and TFE3 returned to baseline levels by 24 hpi (Fig. 2A and 2B).

**Figure 2:**
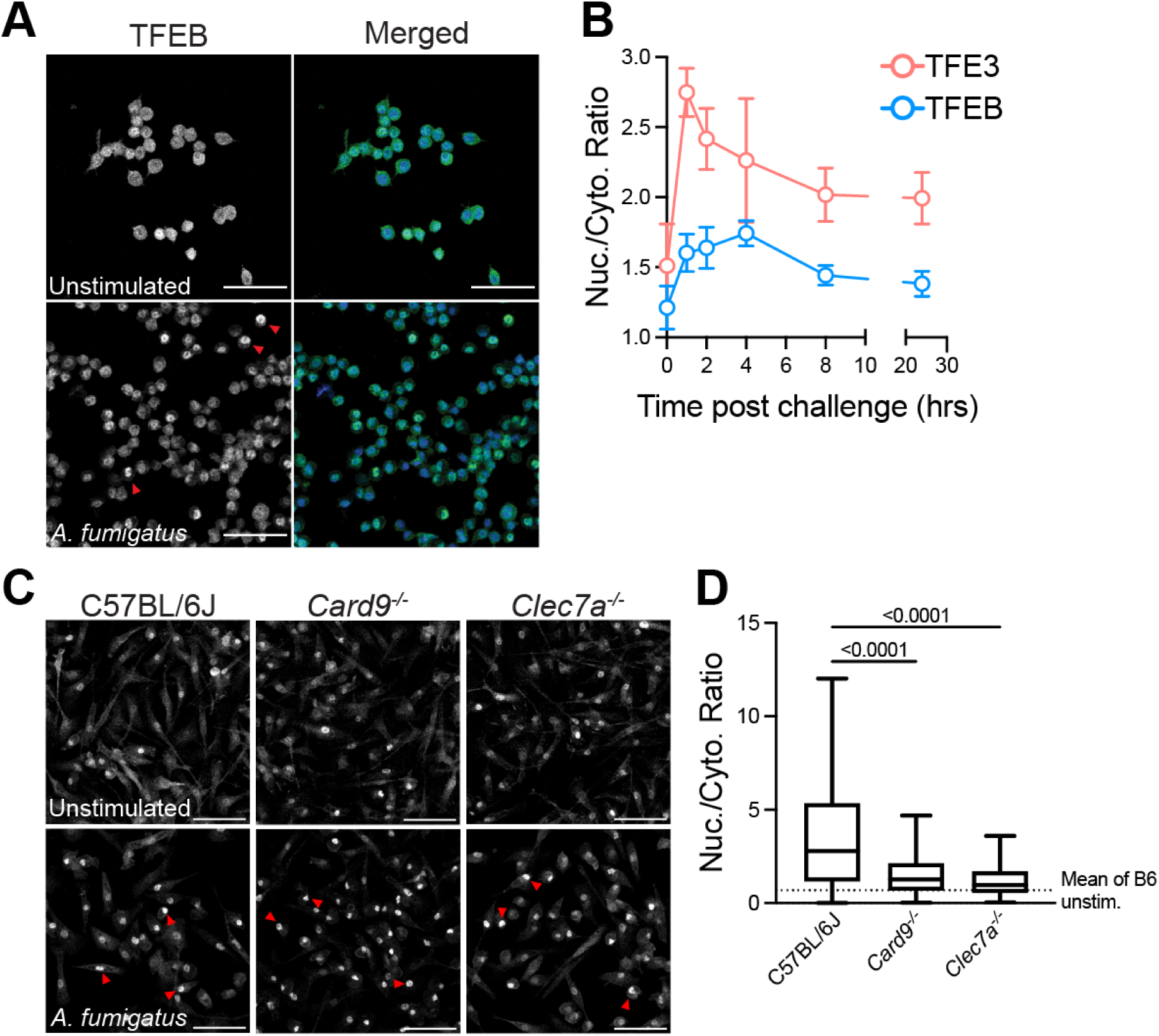
*A. fumigatus* conidia activate TFEB/3 in macrophages via C-type lectin receptors. **(A)** Representative confocal micrographs of iBMDM cells stimulated with swollen heat-killed *A. fumigatus* conidia. Red arrowheads indicate cells with nuclear translocation of TFEB. **(B)** Quantification of TFEB and TFE3 nuclear to cytoplasmic ratio displayed as mean ± SEM from (A). **(C)** Representative confocal micrographs of BMDM cells from B6, *Card9^-/-^*, and *Clec7a^-/-^* mice. Red arrowheads indicate cells with nuclear translocation of TFEB. **(D)** Quantification of TFEB nuclear to cytoplasmic ratio displayed as a box and whiskers plot (C). Scale bar: 50 µm. Data are representative of 2 independent experiments. Statistics: Kruskal-Wallis test and Dunn’s multiple comparisons test.

Phagocytes sense *A. fumigatus* through C-type lectin receptors that bind to fungal cell wall components (e.g., Dectin-1 binds fungal β-glucan), which then signals through spleen tyrosine kinase (Syk) to activate the adaptor protein Card9 to initiate a pro-inflammatory gene expression program (12). We hypothesized that this pathway would also contribute to TFEB and TFE3 activation in response to *A. fumigatus* conidia. To test this hypothesis, we challenged primary BMDMs from *Clec7a^-/-^* (i.e., the gene that encodes Dectin-1) and *Card9^-/-^* mice with *A. fumigatus* conidia and quantified TFEB nuclear translocation. We observed that both *Clec7a^-/-^* and *Card9^-/-^* BMDMs had a reduction in TFEB nuclear translocation following stimulation with *A. fumigatus* conidia compared to WT cells (Fig. 2C and 2D). However, there was residual induction of TFEB nuclear trafficking compared to unstimulated cells, suggesting that additional signals may contribute to TFEB activation in response to *A. fumigatus*. These data suggest that challenge with *A. fumigatus* is sufficient to activate TFEB and TFE3 in phagocytes, and that Dectin-1/CARD9 signaling is required for this process.

### TFEB and TFE3 promote macrophage fungicidal activity *in vitro*

TFEB and TFE3 can drive expression of immune genes in macrophages (19, 22) and activation of TFEB can enhance the bactericidal activity of macrophages (18). Furthermore, lysosomes have been shown to be essential for the control of phagocytosed fungi (20). We hypothesized that TFEB and TFE3 can promote the fungicidal activity of phagocytes. To test this, we generated BMDMs from mice with *Tfeb* deficiency in hematopoietic cells (*Vav-iCre Tfeb^f/f^*; *Tfeb*^Δhem^) since global deletion is embryonic lethal (26). We also generated primary BMDMs from littermates with global *Tfe3* gene deletion (*Tfeb^f/f^ Tfe3^-/y^*; TFE3 KO), or from littermates that lacked *Tfeb* in hematopoietic cells as well as *Tfe3* in all cells (*Vav-iCre Tfeb^f/f^ Tfe3^-/y^*; DKO^hem^). We confirmed successful TFEB depletion in BMDMs by Western blotting (Fig S1A) and in circulating monocytes and neutrophils by flow cytometry (Fig. S1B). BMDMs from DKO^hem^ mice had a reduction in the transcription of two TFEB target genes, *Ctsb* and *Atp6v1c1*, that were upregulated in our lung neutrophil RNA-seq dataset (Fig. S1C), providing further evidence that TFEB and TFE3 were successfully deleted. Next, we challenged BMDMs from DKO^hem^ or *Tfeb^f/f^* mice with FLuorescent *Aspergillus* REporter (FLARE) conidia that are surface-labeled with AF633 and encode RFP, as previously described (27). Upon phagocyte-mediated fungal killing, *Aspergillus* RFP fluorescence is quenched while AF633 fluorescence is preserved, enabling us to quantify the rate of fungal phagocytosis by and fungal viability in diverse phagocyte subsets. The rate of conidial phagocytosis is measured by quantifying total AF633^+^ events for each phagocyte subset analyzed (Uptake = Gate R1 + Gate R2; Fig. 3A). Fungal killing by a specific leukocyte subset is expressed as the frequency of leukocytes that are RFP^+^ and AF633^+^ divided by all AF633+ leukocytes of the same subset independent of RFP fluorescence (i.e., all fungus-engaged leukocytes; Fungal viability in a phagocyte subset = R1/[R1 + R2], Fig. 3A). *In vitro*, we did not observe any difference in the uptake of conidia between DKO and *Tfeb^f/f^* BMDMs (Fig. 3B), but we did observe a significant increase in the fraction of live conidia in DKO compared to *Tfeb^f/f^* BMDMs (Fig. 3C). This finding indicates that genetic loss of TFEB and TFE3 impairs macrophage killing of *A. fumigatus in vitro*.

**Figure 3:**
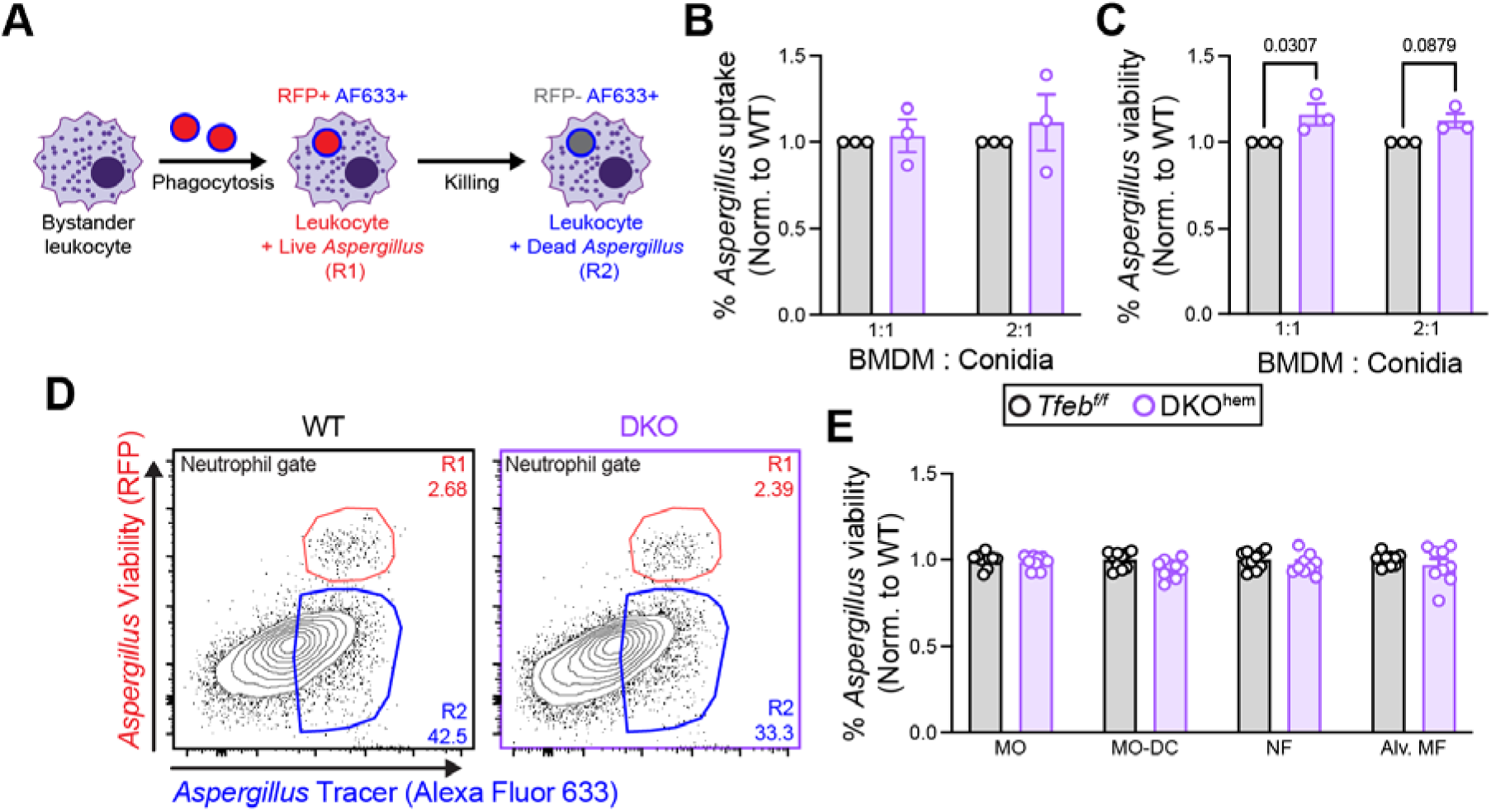
TFEB and TFE3 promote killing of *A. fumigatus in vitro* and *in vivo*. **(A)** Schematic of FLARE strain and changes in fluorescence emission following fungal uptake and killing by host phagocytes. **(B)** The plots show *Aspergillus* uptake (R1 + R2) ± SEM and **(C)** *Aspergillus* viability (R1/(R1 + R2) ± SEM in BMDMs from DKO^hem^ (purple) and *Tfeb^f/f^* (black) after overnight co-culture with FLARE conidia at the indicated ratios. **(D)** Representative plots that display RFP and AF633 fluorescence intensity of lung neutrophils in Tfeb^f/f^ (left panel) or DKO^hem^ (right panel) 24 hpi with 3×10^7^ CEA10 FLARE conidia. **(E)** The plots show conidial viability in alveolar macrophages, neutrophils, monocytes, and Mo-DCs isolated from DKO^hem^ (purple) and *Tfeb^f/f^* (black) 24 hpi with 3×10^7^ FLARE conidia. (B, C, and E) Data from (B and C) 3 or (E) 2 experiments were pooled. (B, C, and E) Dots represent individual mice and data are expressed as mean ± SEM. Statistics: 2-way ANOVA with Sidak’s multiple comparisons test.

Given the defect in fungal killing we observed *in vitro* in DKO macrophages, we hypothesized that TFEB and TFE3 would regulate phagocyte killing in the lung during *A. fumigatus* infection. To test this, we challenged DKO or *Tfeb^f/f^* mice with FLARE conidia, and 24 hpi, we harvested the lungs and quantified the phagocytosis and killing of *A. fumigatus* by lung phagocytes as well as phagocyte numbers in the lung (Fig. S2A). DKO mice had slightly greater numbers of lung monocytes and Mo-DCs but had no difference in alveolar macrophages or neutrophils compared to control mice (Fig. S2B). We observed no difference in the phagocytosis of *A. fumigatus* conidia by lung phagocytes in DKO mice compared to *Tfeb^f/f^* (Fig. S2C), consistent with our *in vitro* findings (Fig. 3B left). In contrast to *in vitro* results with BMDMs, DKO lung phagocytes exhibited no defect in conidial killing (Fig. 3D and 3E) since the frequency of live fungal cells was similar in DKO and control leukocytes for each phagocyte population analyzed. This result suggests that while TFEB and TFE3 promote the fungicidal activity of macrophages *in vitro*, their expression is dispensable for fungal killing by myeloid phagocytes *in vivo*.

To examine whether loss of TFEB and TFE3 would abrogate the increase in LysoTracker signal that we observed in phagocytes upon fungal uptake (Fig. 1D), we infected *Tfeb^f/f^* mice and DKO^hem^ mice with *A. fumigatus*, and measured LysoTracker staining in lung single cell populations by flow cytometry at 24 hpi. Fungus-engaged phagocytes isolated from DKO^hem^ mice and from *Tfeb^f/f^* control mice showed an increase in LysoTracker staining compared to bystander phagocytes. Thus, expression of TFEB and TFE3 was not required for the increase in LysoTracker staining observed with fungal cell uptake (Fig. S3A and S3B in the supplemental material). These data support the notion that loss of TFEB and TFE3 does not impair lysosomal acidification in response to fungal engagement *in vivo*, which may explain why myeloid phagocytes do not exhibit an impairment in fungal killing.

### TFEB and TFE3 are dispensable for immunity to *A. fumigatus* in the lung

While the contributions of TFEB and TFE3 to immune function have been extensively studied *in vitro* and in non-vertebrate animal models (22, 28), data on the role of TFEB and TFE3 in experimental murine infection models, and in the context of fungal infections, are lacking. To test the requirement for TFEB and TFE3 in hematopoietic cells in host defense against *A. fumigatus,* we generated bone marrow chimeric mice, using *Tfeb*^Δhem^, DKO^hem^ or *Tfeb^f/f^* donor bone marrow injected intravenously into irradiated recipient mice. Bone marrow chimeric mice were rested for 6 weeks and subsequently challenged mice with *A. fumigatus* conidia via the intratracheal route. We also challenged global TFE3 KO and *Tfeb^f/f^*control mice in separate experiments. All genotypes of mice were resistant to infection with *A. fumigatus* (Fig. 4A-C). We did not observe a significant difference in *A. fumigatus* lung fungal burden in any group at 24 hpi (Fig. 4(D-F). These findings indicate that TFEB (in hematopoietic cells) and TFE3 (globally) are not essential for clearance of *Aspergillus* from the lung or for survival following infection.

**Figure 4.**
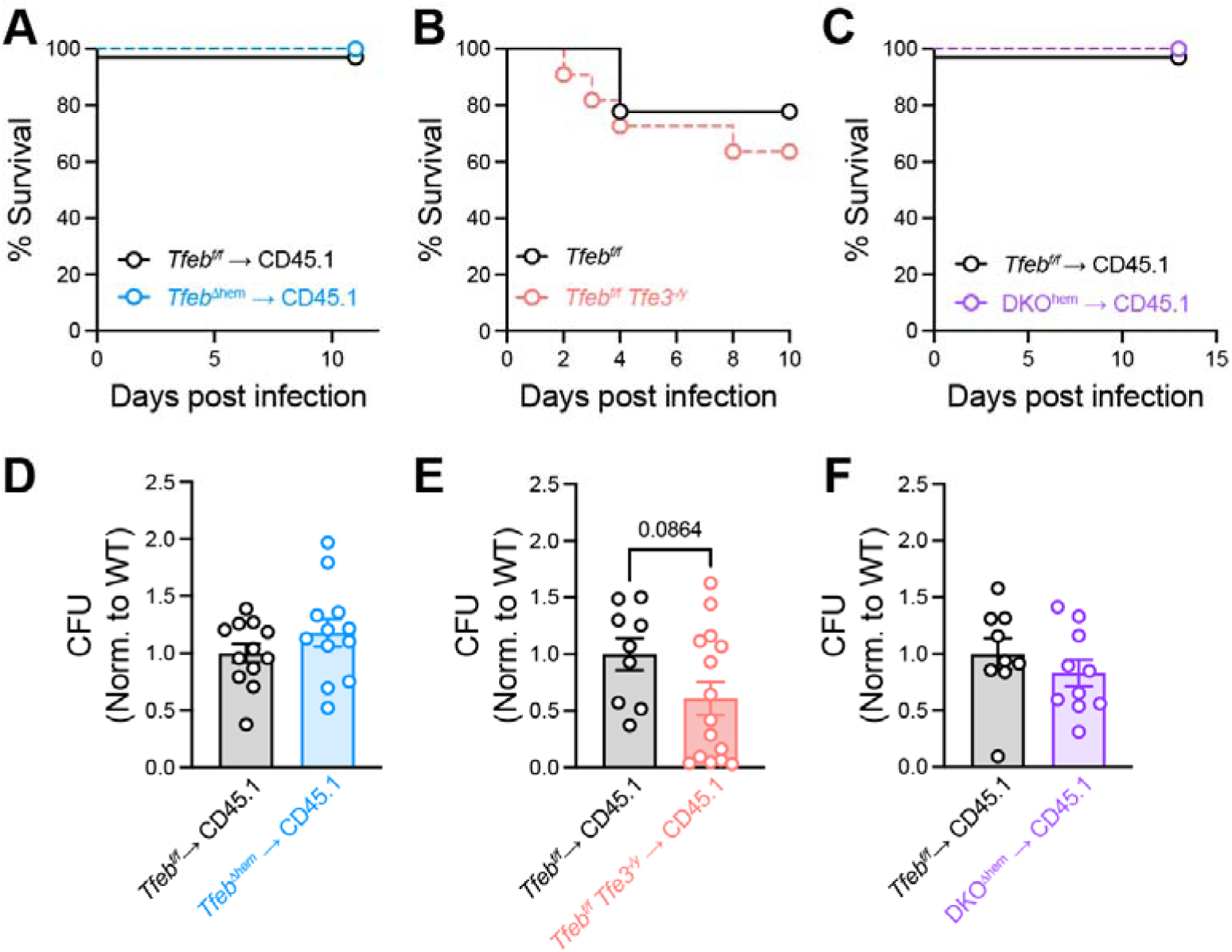
TFEB/3 are dispensable for murine survival during *A. fumigatus* infection. **(A-C)** Kaplan-Meier survival curve of (A) *Tfeb^f/f^*→ CD45.1 (n = 7) and *Tfeb*^Δhem^ → CD45.1 (n = 8), (B) *Tfebf/f* (n = 10) and TFE3 KO (n = 11), and (C) *Tfeb^f/f^*→ CD45.1 (n = 4) and DKO^hem^ → CD45.1 (n = 6) hematopoietic chimera mice challenged with 6×10^7^ CEA10 conidia. **(D-F)** Lung fungal burden at 24 h.p.i. with 3×10^7^ FLARE conidia in *Tfeb^f/f^*→ CD45.1, *Tfeb*^Δhem^ → CD45.1, *Tfeb^f/f^ Tfe3^-/y^* → CD45.1, and DKO^hem^ → CD45.1. (D-F) Data are pooled from 2 independent experiments and each dot indicates a mouse. Statistics: Mann-Whitney test.

## DISCUSSION

This study aimed to investigate the role of transcription factors TFEB and TFE3 in phagocyte function during immune responses to *A. fumigatus*. We found that lysosome-related genes are upregulated in lung neutrophils following fungal engagement, and that *A. fumigatus* conidia activate TFEB and TFE3 myeloid phagocytes, as determined by nuclear translocation. Activation of TFEB and TFE3 in response to *A. fumigatus* was impaired by deletion of Dectin-1 and CARD9, linking *Aspergillus* recognition to TFEB and TFE3 activation. Additionally, *Tfeb* and *Tfe3* deletion in macrophages led to a reduction in fungicidal activity *in vitro*, although in the lung, no impairment in killing was observed. Interestingly, mice with hematopoietic TFEB deletion, TFE3 deletion, or combined deletion in the hematopoietic compartments were resistant to infection with *A. fumigatus*, indicating functional redundancy. Overall, this study provides valuable insights into the activation of TFEB and TFE3 in response to *A. fumigatus* conidia and the contribution of lysosome expansion to the effector functions of phagocytes upon fungal recognition. The study also adds to the growing literature on the role of TFEB and TFE3 in antifungal and antibacterial immunity.

Innate immune cells are known to employ various regulatory mechanisms upstream of TFEB and TFE3 to respond to pathogens. Phagocytosis of IgG-opsonized latex beads activates TFEB in murine RAW264.7 cells and primary BMDMs via FcγR and Syk (18). Syk is also a critical component of C-type lectin receptor signaling, linking Dectin-1 to Card9 (29). In response to *A. fumigatus*, Dectin-1 and Card9 are required for optimal activation of TFEB and TFE3, suggesting that Syk may also be involved in this context. Several groups have shown that calcium flux from the lysosome via MCOLN1 and activation of calcineurin are critical for activation of TFEB and TFE3 in macrophages (18, 19) and other cell types (30). Fungal pathogen sensing via C-type lectin receptors can also trigger calcium flux in phagocytes (31). While we did not formally test the requirement for calcium and calcineurin, it is possible that these molecules are also important for TFEB and TFE3 activation in response to *A. fumigatus*. Inhibition of mTOR also activates TFEB and TFE3 (32–34). However, it was recently shown that TFEB could be active in the context of hyperactive mTORC1 due to unique interactions between TFEB and Rag GTPases, suggesting that in specific settings, mTOR inhibition is not strictly required for TFEB and TFE3 activation (35). Since *A. fumigatus* is known to activate mTORC1 signaling in macrophages (36), fungus-induced TFEB and TFE3 activation likely occurs in the presence of active mTORC1.

TFEB and TFE3 play important roles in the regulation of lysosome biogenesis and autophagy in various cell types, including macrophages (28). While overexpression of TFEB was found to increase lysosome biogenesis in murine macrophages (37), we did not observe significant differences in LysoTracker staining between TFEB and TFE3 knockout cells and wild-type cells, suggesting functional redundancy in lysosomal biogenesis and acidification in phagocytes during *A. fumigatus* infection. TFEB was also identified as a critical mediator of autophagy in *Mycobacterium tuberculosis*-infected macrophages (38). *A. fumigatus-*containing phagosomes can recruit LC3 and deletion of *Atg5* in hematopoietic cells renders mice susceptible to *A. fumigatus* in a corticosteroid model of infection (39), consistent with an anti-*Aspergillus* role for autophagy pathway proteins in pharmacologically immune compromised hosts. We did not examine LC3 or *Atg5* induction in our BMDM or *in vivo* experiments, but the resistance observed in DKO mice to *A. fumigatus* infection suggests that regulation of the autophagy pathway may be controlled by independent mechanisms. Notably, our study differs from the previous investigation in which corticosteroids were used as part of the *A. fumigatus* infection model in Atg5-deficient mice. We performed studies in gene knockout mice without pharmacologic immune suppression, and this difference in methodology may explain the divergent results between our study and theirs.

TFEB and TFE3 were found to directly control the expression of several cytokines and chemokines in murine macrophages infected with pathogenic bacteria or stimulated with LPS (18, 21). Given the lack of fungal clearance defect or differences in survival in mice that lack TFEB and TFE3 expression in hematopoietic cells, we did not measure TFEB- or TFE3-dependent lung cytokine levels. It is thus unlikely that TFEB or TFE3 regulate the levels of essential mediators of intercellular crosstalk and host defense in the lung (e.g., GM-CSF (40), type I and type III IFN (41), or CXCL9 and CXCL10 (42)). It is possible that cytokine-independent compensatory mechanisms exist to protect TFEB- and TFE3-deficient mice during infection, such as greater leukocyte recruitment (Fig. S2) or increased fungicidal activity via NADPH oxidase.

In this study, we found that during *A. fumigatus* infection, phagocytes activate TFEB and TFE3 and expand their lysosomal compartment. While this pathway is activated *in vitro* and *in vivo* and is essential for optimal *A. fumigatus* conidiacidal activity *in vitro*, TFEB and TFE3 are ultimately not essential for murine survival during infection and for phagocyte function *in vivo*. This work contributes to our understanding of pathways that are activated in phagocytes upon *A. fumigatus* challenge and how lysosomes and TFEB/TFE3 contribute to phagocyte function.

## ACKNOWLEDGMENTS

We thank all members of the Hohl laboratory for insightful discussions. We thank Anupam Jhingran for help with generating the RNA-sequencing data. We thank Eric Chan for help with microscope operation and image analysis. The studies were supported by NIH grants P30 CA008748 (to MSKCC), R37 AI093808 (TMH), R01 AI139632 (TMH), F31 AI161996 (MAA), and F31 AI167511 (KAMM). The funders had no role in study design, data collection and analysis, decision to publish or preparation of manuscript. We acknowledge assistance from the MSKCC core facilities including Bioinformatics Core, Flow Cytometry Core Facility, Integrated Genomics Operation, Molecular Cytology Core Facility.

## AUTHOR CONTRIBUTIONS

Conceptualization, T.M.H., M.A.A; Methodology, T.M.H.; Investigation, M.A.A., K.A.M.M, M.G., N.S.; Writing – Original Draft, M.A.A. and T.M.H.; Writing – Review & Editing, M.A.A. and T.M.H.; Funding Acquisition, T.M.H. and M.A.A; Resources, A.B.

## DECLARATION OF INTERESTS

A. Ballabio is co-founder of CASMA Therapeutics and advisory board member of Next Generation Diagnostics and Avilar and Coave Therapeutics. All other authors declare no competing interests.

## METHODS

### Mice

C57BL/6J mice (cat#: 000664) and *Vav-iCre* mice (cat#: 008610) were purchased from The Jackson Laboratory. CD45.1^+^ C57BL6.SJL mice (cat#: 556) were purchased from Charles River Laboratories. *Tfeb^f/f^*mice (34) were crossed with *Vav-iCre* mice to generate *Tfeb*^Δ*hem*^. *Tfeb*^Δ*hem*^ mice were further crossed to *Tfe3^-/y^*mice(43) to generate *Tfeb*^Δ*hem*^ *Tfe3^-/y^* (DKO) mice. *Card9^-/-^* mice (44) were provided by Dr. Xin Lin (Tsinghua University). *Clec7a^-/-^* mice (45) were provided by Dr. Shinobu Saijo (University of Tokyo). All mouse strains were bred and housed in the MSKCC Research Animal Resource Center under specific pathogen-free conditions. All animal experiments were conducted with sex- and age-matched mice and performed with MSKCC Institutional Animal Care and Use Committee approval. Wherever possible, littermate controls were utilized. Animal studies complied with all applicable provisions established by the Animal Welfare Act and the Public Health Services Policy on the Humane Care and Use of Laboratory Animals.

### *Aspergillus fumigatus* culture and Murine Infection Model

*A. fumigatus* CEA10 (46) and CEA10-RFP (47) strains were cultured on glucose minimal medium slants at 37 °C for 4-7 days prior to harvesting conidia for experimental use. To generate AF633-labeled or FLARE conidia for experimental use, 7×10^8^ CEA10 (for AF633-labeled) or CEA10-RFP (for FLARE) conidia were rotated in 10 μg/ml Sulfo-NHS-LC-Biotin (Thermo Scientific) in 1 ml of 50 mM carbonate buffer (pH 8.3) for 2 hr at 4 °C, incubated with 20 μg/ml Streptavidin, Alexa Fluor™ 633 conjugate (Molecular Probes) at 37 °C for 1 h, resuspended in PBS and 0.025% Tween 20 for use within 24 hr (12).

To generate morphologically uniform heat-killed swollen conidia, 5×10^6^/ml conidia were incubated at 37 °C for 14 hours in RPMI-1640 and 0.5 μg/ml voriconazole and heat-killed at 100 °C for 30 minutes (48). To infect mice with 3-6×10^7^ *A. fumigatus* cells, conidia were resuspended in PBS, 0.025% Tween-20 at a concentration of 0.6-1.2×10^9^ cells/mL and 50 µl of cell suspension was administered via the intratracheal route to mice anesthetized by isoflurane inhalation.

### Transcriptomic analysis

Bystander and fungus-engaged neutrophils were sorted from the lungs of mice infected with AF633-labeled *A. fumigatus* 24 hpi. Dead cells were excluded using DAPI (ThermoFisher cat#: EN62248). Bystander neutrophils were purified (>95% purity) by sorting as CD45^+^, Ly6G^+^, Ly6B^+^, CD11b^+^, and AF633^-^ cells, and fungus-engaged neutrophils were purified (>95%) by sorting as CD45^+^Ly6G^+^, Ly6B^+^, CD11b^+^, and AF633^+^ cells using a BD FACSAria cell sorter.

For RNA extraction sorted cells were lysed in TRIzol Reagent (ThermoFisher catalog # 15596018) and phase separation was induced with chloroform. Neutrophil-derived RNA was precipitated with isopropanol and linear acrylamide, washed with 75% ethanol, and resuspended in 15 µL nuclease-free water. For transcriptome sequencing, after RiboGreen quantification and quality control by Agilent BioAnalyzer, 10 ng total RNA with RNA integrity numbers ranging from 8.8 to 9.5 underwent amplification using the SMART-Seq v4 Ultra Low Input RNA Kit (Clonetech catalog # 63488), with 12 cycles of amplification. Subsequently, 10 ng of amplified cDNA was used to prepare libraries with the KAPA Hyper Prep Kit (Kapa Biosystems KK8504) using 8 cycles of PCR. Samples were barcoded and run on a HiSeq 2500 in a PE50 run, using the TruSeq SBS Kit v4 (Illumina). An average of 42 million paired reads were generated per sample and the percent of mRNA bases per sample ranged from 73% to 84%.

Statistical analysis of RNASeq data was performed by the MSKCC Bioinformatics Core. The output data (FASTQ files) were mapped to the target genome using the rnaStar aligner that maps reads on a genomic basis and resolves reads across splice junctions. The 2pass mapping method [103] was used, in which the first mapping pass uses a list of known annotated junctions from Ensembl. Novel junctions found in the first pass were then added to the known junctions and a second mapping pass was performed in which the Remove Noncanonical flag was used. After mapping, the output SAM files were post processed using the PICARD tool AddOrReplaceReadGroups to add read groups, sort the files, and covert them to the compressed BAM format. The expression count matrix was then computed from the mapped reads using HTSeq (www-huber.embl.de/users/anders/HTSeq) and one of several possible gene model databases. The raw count matrix generated by HTSeq was then processed using the R/Bioconductor package DESeq (www-huber.embl.de/users/anders/DESeq) which was used to both normalize the full dataset and analyze differential expression between sample groups.

### LysoTracker staining

Total intracellular lysosomal content was measured using LysoTracker Green DND-26 (cat#: L7526). Briefly, single cell lung suspensions were incubated in complete DMEM with 50 nM LysoTracker at 37 °C for 45 min according to manufacturer’s instruction. Cells were then stained with antibodies against surface markers as previously described and analyzed by flow cytometry.

### Immunofluorescent staining and imaging of TFEB and TFE3

Immortalized bone marrow derived macrophages (iBMDMs) (49) were cultured in complete DMEM. Cells were passaged 1:10 every 3 days. To assess the subcellular localization of TFEB or TFE3 in iBMDM cells, 5×10^5^ cells were seeded into 12 well plates on top of coverslips in cDMEM. Swollen heat killed conidia (SHKC) were prepared as described above and 5×10^6^ SHKC were added to the appropriate wells (MOI = 10). Following stimulation, cells were fixed using 4% paraformaldehyde (PFA) in PBS and permeabilized with PBS containing 0.25% Triton X-100. Cells were then incubated with rabbit α-TFEB (1:1000) (ThermoFisher cat#: 50-156-5746) or rabbit α-TFE3 (1:1000) (ThermoFisher cat#: HPA023881) followed by incubation with goat anti-rabbit IgG AF488 (ThermoFisher cat#: A-11008). Finally, cells were counterstained with 1 µg/mL DAPI, mounted on slides with Mowiol 4-88 (ThermoFisher cat#: 47-590-4100GM), and imaged using a Leica TCS SP5 Confocal Microscope. Nuclear TFEB or TFE3 were calculated by outlining DAPI-positive structures and calculating the average TFEB or TFE3 staining intensity in each region, using Fiji (50).

### BMDM differentiation

Tibias and femurs were removed from euthanized mice of the indicated genotype, cut with scissors at each end, and centrifuged to harvest bone marrow cells, as previously described (51). Bone marrow cells underwent red blood cell lysis and were plated at 1×10^7^ bone marrow cells per 10 cm dish in macrophage differentiation media consisting of DMEM + 20% Fetal Bovine Serum + 30% L-929 M-CSF conditioned + 2 mM L-glutamine + Pen/Strep (all purchased from ThermoScientific). Plates were fed with 5 mL of additional macrophage differentiation media on day 3 of differentiation. Cell culture media was removed on day 7 and day 8 cultures. Cells were detached from non-treated plates with cold RPMI + 3 mM EDTA. To analyze conidia uptake and killing, FLARE conidia were co-cultured with BMDMs at the indicated ratios overnight in DMEM containing 10% FBS, Penicillin and Streptomycin (Pen+Strep), and L-glutamine and HEPES. This media is referred to below as complete DMEM. For co-culture with *A. fumigatus*, 0.5 μg/ml voriconazole was added to the media. BMDMs were detached with cold RPMI containing 3 mM EDTA, transferred to a 96 well U-bottom plate, and washed twice with PBS + 0.5% bovine serum albumin (ThermoScientific cat#: BP9703100) + 0.5 mM EDTA. DAPI (0.1 µg/mL) was added to cells immediately before being analyzed by flow cytometry.

### Flow cytometric analysis of BMDMs and single cell lung suspensions

Single cell lung suspensions were prepared for flow cytometry as described in (52), with minor modifications. Briefly, perfused murine lungs were placed in a gentle MACS C tube and mechanically homogenized in 5 ml of PBS using a gentle MACS Octo Dissociator (Miltenyi Biotec). Lung cell suspensions were lysed of RBCs, enumerated, and stained with fluorophore-conjugated antibodies prior to flow cytometric analysis on a Beckman Coulter Cytoflex LX and analyzed with FlowJo version 10.8.1. Dead cells were excluded using LIVE/DEAD Fixable Aqua Dead Cell Stain (ThermoFisher cat#: L34957). The antibodies used are the following: anti-Ly6C (clone AL-21), anti-Ly6G (clone 1A8), anti-CD11b (clone M1/70), anti-CD11c (clone HL3), anti-CD45 (clone 30-F11), anti-I-A/I-E (clone M5/114.15.2), anti-Ly6B.2 (clone 7/4), and anti-Siglec-F (clone E50-2440) all from Biolegend and BD Biosciences. Alveolar macrophages were identified as CD45^+^ CD11c^+^ Siglec-F^+^ cells, neutrophils were identified as CD45^+^ CD11b^+^ Ly6G^+^ cells, inflammatory monocytes as CD45^+^ CD11b^+^ CD11c^-^ Ly6G^-^ Ly6C^hi^ cells, and Mo-DCs as CD45^+^ CD11b^+^ CD11c^+^ Ly6G^-^ Ly6C^hi^ MHC-II^+^ cells. To analyze the lung fungal burden, perfused murine lungs were homogenized using a PowerGen 125 homogenizer (Fisher) in 2 mL PBS, 0.025% Tween-20, and plated on Sabouraud dextrose agar.

In data analyses for a given leukocyte subset, conidial uptake refers to the frequency of fungus-engaged leukocytes i.e., the sum of RFP^+^AF633^+^ and RFP^-^AF633^+^ leukocytes. Conidial viability within a specific leukocyte subset refers to the frequency of leukocytes that contain live conidia (RFP^+^AF633^+^) divided by the frequency of all fungus-engaged leukocytes (RFP^+^AF633^+^ and mCherry^-^AF633^+^).

### Generation of bone marrow chimeric mice

For single bone marrow chimeras, CD45.1^+^ C57BL6.SJL recipients were lethally irradiated (900cG), reconstituted with either 2-5×10^6^ CD45.2^+^ *Tfeb^f/f^*, *Tfeb^f/f^ Tfe3^-/y^*, *Vav^iCre/+^ Tfeb^f/f^*, or *Vav^iCre/+^ Tfeb^f/f^ Tfe3^-/y^* donor bone marrow cells isolated from littermates. After bone marrow transplantation, recipient mice received 400 μg/ml enrofloxacin in the drinking water for 21 days to prevent bacterial infections and rested for 6-8 weeks prior to experimental use.

### Western blotting

BMDMs were lysed in lysis buffer (150 mM NaCl, 50 mM HEPES pH 7.4, 1 mM EDTA, 1% Nonidet P-40, protease inhibitors). Total cell lysates were subjected to SDS–PAGE and then blotted with the indicated antibodies.

### Quantitative PCR Analysis

For quantitative RT-PCR, BMDMs were stimulated with *A. fumigatus*, and RNA was extracted using TRIzol Reagent. cDNA was generated from RNA using the Applied Biosystems High-Capacity RNA-to-cDNA Kit (ThermoFisher cat#: 4387406), and qRT-PCR was performed on aStepOnePlus Real Time PCR System (Applied Biosystems) using TaqMan Fast Advanced Master Mix and TaqMan Gene Expression Assays (ThermoFisher Scientific) *Atp6v1c1* (Mm00445925) and *Ctsb* (Mm01310506).

### Quantification and statistical analysis

Data are representative of at least 2 independent experiments, as indicated. All results are expressed as mean (± SEM), unless stated otherwise. The Mann-Whitney test was used for comparisons of two groups, unless noted otherwise. The Kruskal-Wallis test was used for multi-group comparisons, unless noted otherwise. Survival data was analyzed by log-rank test. All statistical analyses were performed with GraphPad Prism software, v8.2.0.

### Resource Availability

Further information and requests for resources or reagents should be directed to the Lead Contact, Tobias M. Hohl.

### Materials Availability

The datasets generated for this study can be found in NCBI GEO, NCBI accession no. GSE233353.

## SUPPLEMENTAL FIGURES

**Supplemental Figure 1:**
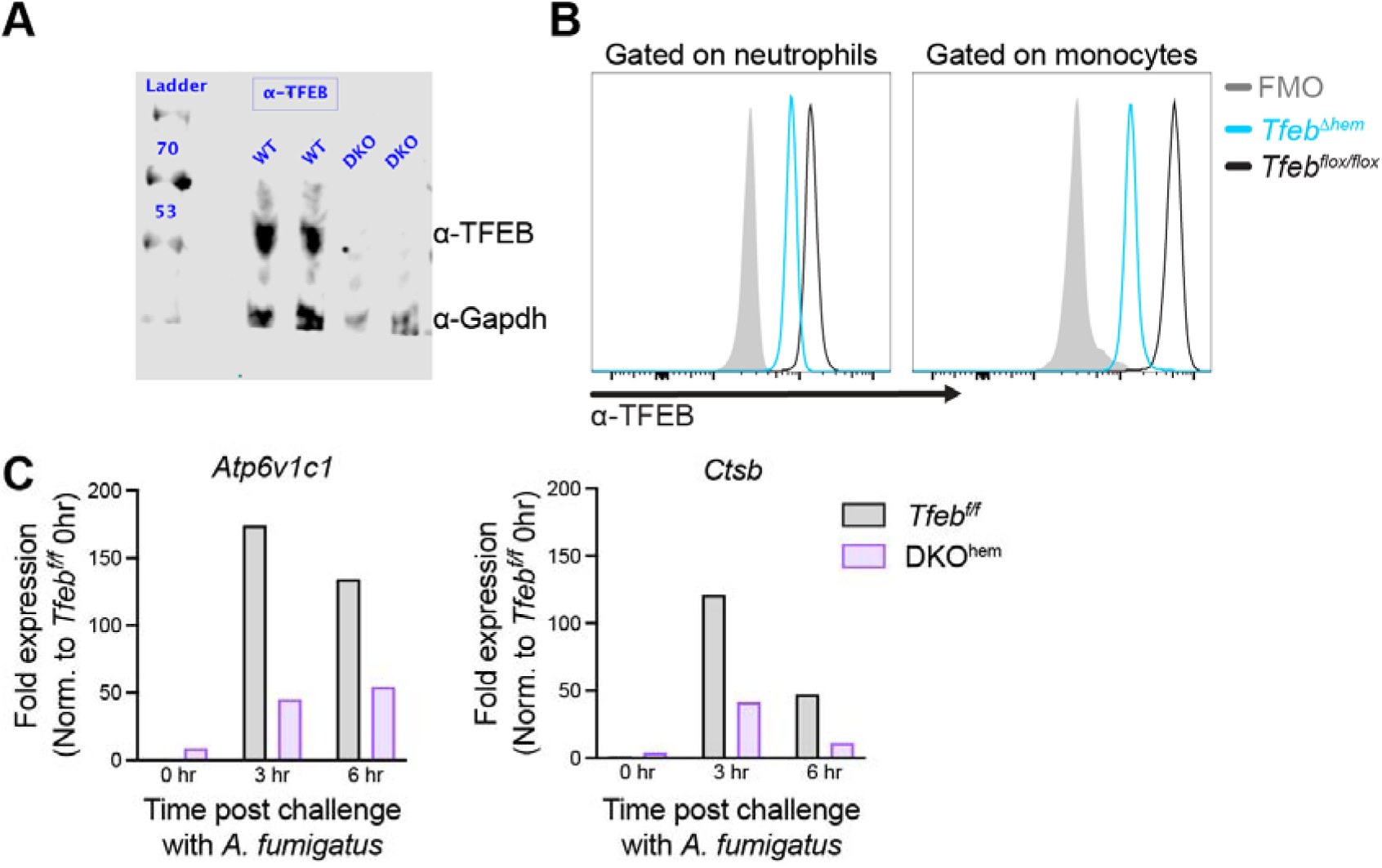
Deletion of TFEB in hematopoietic cells using Vav-iCre. (A) Western blots of TFEB (left panel) and TFE3 (right panel) from BMDMs differentiated from *Tfeb^f/f^* mice and DKO^hem^ mice. (B) Quantitative PCR of TFEB and TFE3 target genes in *Tfeb^f/f^* and DKO BMDMs following stimulation with *A. fumigatus* swollen heat-killed conidia. Data are from 1 experiment performed in duplicate (A), or triplicate (B and C).

**Supplemental Figure 2:**
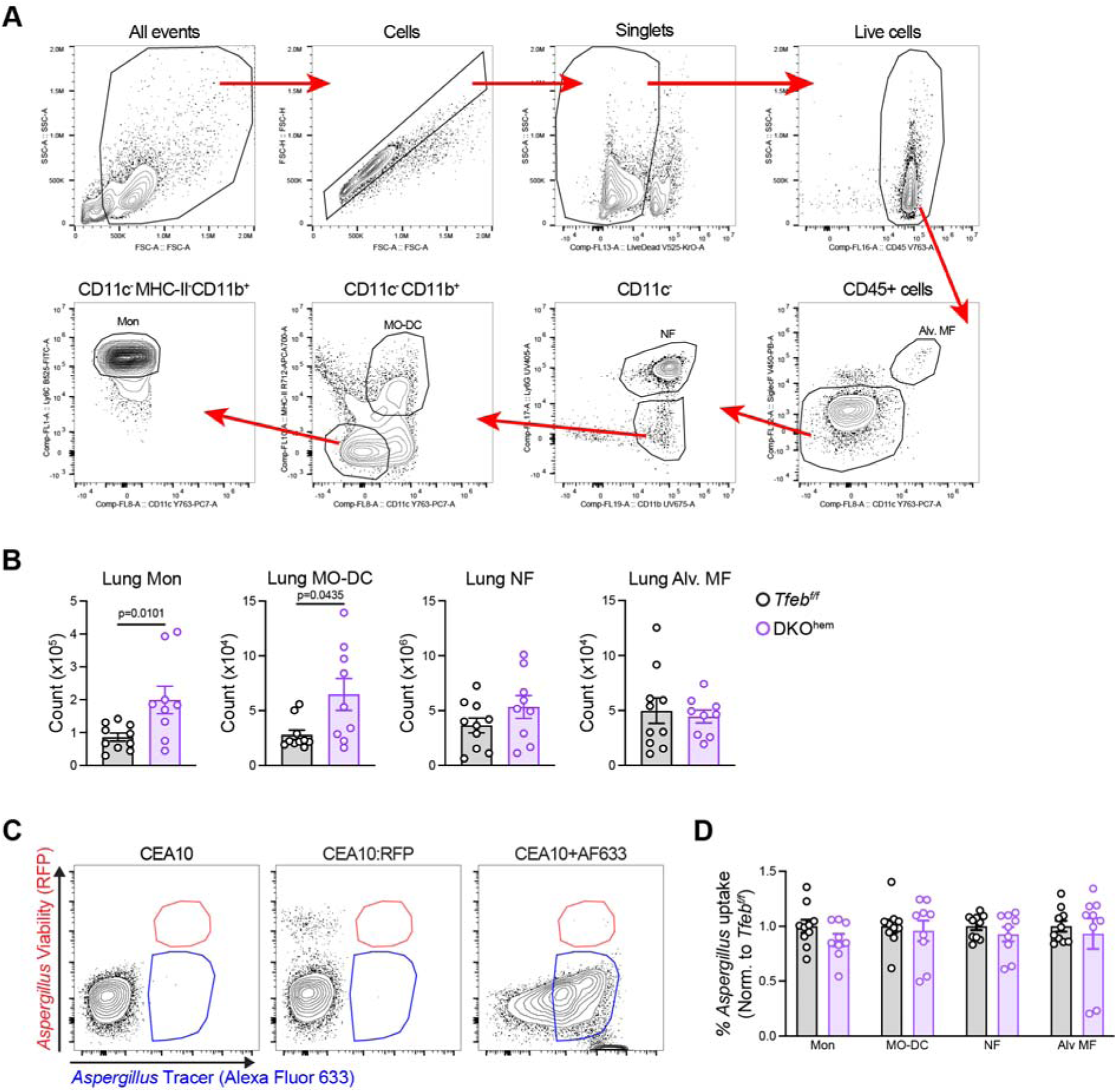
Gating strategy for lung FLARE experiments. (A) Gating strategy to identify lung phagocyte subsets at 24 hpi. (B) The plots indicate the number of lung phagocytes at 24 hpi from mice of the indicated genotypes (need legend). (C) Representative flow cytometry plots of fluorescence minus one control (FMOs) used to gate R1 and R2 cells. (D) The plots indicate *Aspergillus* uptake by lung phagocytes at 24 hpi normalized to *Tfeb^f/f^* mice. Data from 2 experiments were pooled. (B and D) Dots represent individual mice and data are expressed as mean ± SEM. Statistics: (B) Mann-Whitney test.

**Supplemental Figure 3:**
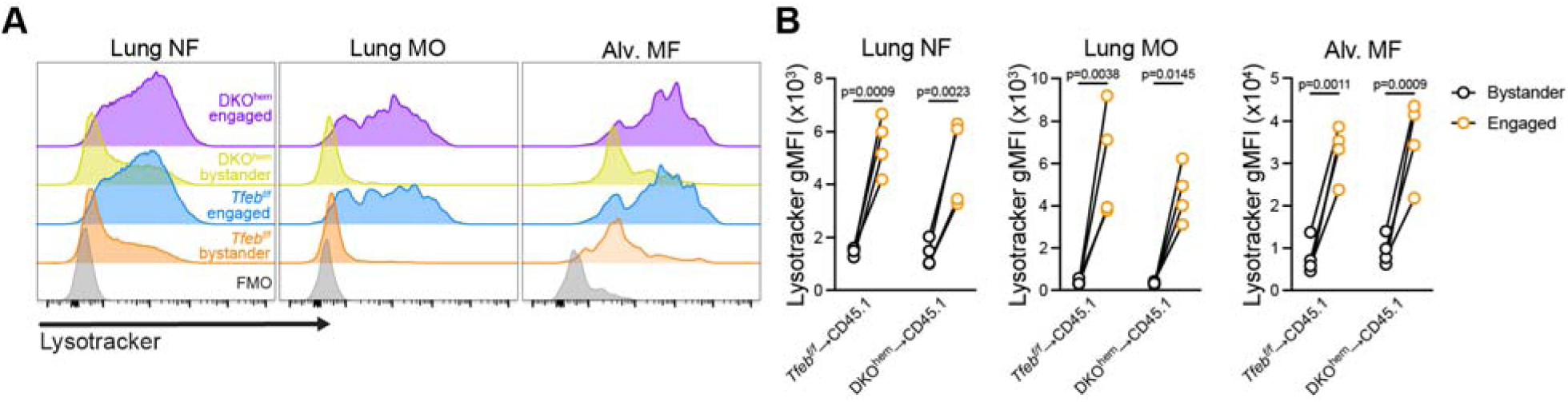
LysoTracker staining in lung phagocytes from DKO^hem^ mice during *A. fumigatus* infection. (A) Representative histograms of LysoTracker signal in lung phagocytes 24 hpi with *A. fumigatus*. (B) Quantification of LysoTracker gMFI. Dots represent individual mice with lines depicting results obtained from bystander and fungus-engaged cells from the same mouse. Statistics: RM two-way ANOVA with Sidak’s multiple comparisons test.

**Supplemental table 1:** TFEB target genes upregulated in engaged lung neutrophils.

|  |
| --- |
| Atp6v0d2 |
| Atp6v1a |
| Atp6v1c1 |
| Cd63 |
| Clcn7 |
| Ctns |
| Ctsb |
| Gaa |
| Gla |
| Hexa |
| Lamp1 |
| Naglu |
| Plbd2 |
| Tpp1 |
| Vps33a |

